# Overdosage of balanced protein complexes reduces proliferation rate in aneuploid cells

**DOI:** 10.1101/376988

**Authors:** Ying Chen, Siyu Chen, Ke Li, Yuliang Zhang, Xiahe Huang, Ting Li, Shaohuan Wu, Yingchun Wang, Lucas B. Carey, Wenfeng Qian

**Affiliations:** State Key Laboratory of Plant Genomics, Institute of Genetics and Developmental Biology, Chinese Academy of Sciences, Beijing 100101, China; State Key Laboratory of Molecular Developmental Biology, Institute of Genetics and Developmental Biology, Chinese Academy of Sciences, Beijing 100101, China; Key Laboratory of Genetic Network Biology, Institute of Genetics and Developmental Biology, Chinese Academy of Sciences, Beijing 100101, China; University of Chinese Academy of Sciences, Beijing 100049, China; Department of Experimental and Health Sciences, Universitat Pompeu Fabra, Barcelona, 08003, Spain

**Keywords:** Aneuploidy, protein complexes, cell proliferation, yeast

## Abstract

Cells with complex aneuploidies, such as tumor cells, display a wide range of phenotypic abnormalities. However, molecular basis for this has been mainly studied in trisomic (2*n*+1) and disomic (*n*+1) cells. To determine how karyotype affects proliferation rate in cells with complex aneuploidies we generated forty 2*n+x* yeast strains in which each diploid cell has an extra 5 to 12 chromosomes and found that these strains exhibited abnormal cell-cycle progression. Proliferation rate was negatively correlated with the number of protein complexes in which all subunits were at the 3-copy level, but not with the number of imbalanced complexes made up of a mixture of 2-copy and 3-copy genes. Proteomics revealed that most 3-copy members of imbalanced complexes were expressed at only 2*n* protein levels whereas members of complexes in which all subunits are stoichiometrically balanced at 3 copies per cell had 3n protein levels. We identified individual protein complexes for which overdosage reduces proliferation rate, and found that deleting one copy of each member partially restored proliferation rate in cells with complex aneuploidies. Lastly, we validated this finding using orthogonal datasets from both yeast and from human cancers. Taken together, our study provides a novel explanation how aneuploidy affects phenotype.

## INTRODUCTION

Aneuploidy refers to karyotypes that are not an exact multiple of the haploid genome (Birchler and Veitia, 2007; Täckholm, 1922; Torres et al., 2008; Torres et al., 2010b) and is a hallmark of tumor cells (Albertson et al., 2003; Holland and Cleveland, 2009; Siegel and Amon, 2012; Weaver and Cleveland, 2006). At the cellular level, aneuploidy usually exhibits a reduced proliferation rate (Birchler and Veitia, 2012; Torres et al., 2008). For example, in the budding yeast *Saccharomyces cerevisiae*, proliferation rate of aneuploidy is reduced significantly under non-stress conditions (Parry and Cox, 1970; Pavelka et al., 2010). Similar observations were also reported in the fission yeast *Schizosaccharomyces pombe* (Niwa et al., 2006; Niwa and Yanagida, 1985) and mammalian cells (Baker et al., 2004; Segal and McCoy, 1974; Stingele et al., 2012; Thompson and Compton, 2008; Williams et al., 2008). Aneuploidy has also been suggested to suppress the proliferation of tumor cells (Sheltzer et al., 2017). The apparent fast expansion of tumor cell population is likely because mutations in cell cycle-related genes remove the proliferative restrictions imposed on individual cells in multicellular organisms, obscuring the adverse effect of aneuploidy (Holland and Cleveland, 2009; Torres et al., 2008; Torres et al., 2010b).

Although the reduced proliferation rate has been observed in many types of aneuploid cells, the mechanistic link between the two remains under investigation. Two mutually non-exclusive hypotheses have been proposed. The balance hypothesis asserts that the disruption of the stoichiometric relationship among subunits in a macromolecular complex perturbs its function and can cause cytotoxicity (Birchler and Veitia, 2007, 2010; Papp et al., 2003; Veitia, 2002, 2005). Consistent with the balance hypothesis, gain or loss of some chromosomes in a genome usually results in greater growth defects than complete duplication of all chromosomes (Birchler and Veitia, 2010; Otto and Whitton, 2000). In addition, the overexpression of the α-tubulin gene can partly rescue the lethality caused by the overexpression of the β-tubulin gene (Abruzzi et al., 2002; Katz et al., 1990), demonstrating the importance of dosage balance within a protein complex. The burden hypothesis was proposed based on the observation that aneuploid cells were under proteotoxic stress (Oromendia et al., 2012), which implies that the synthesis, folding, and degradation of the extra protein subunits are a burden to aneuploid cells (Dephoure et al., 2014; Torres et al., 2010b). In fact, 50% to 70% of extra subunits in an imbalanced complex are degraded (Dephoure et al., 2014; Ishikawa et al., 2017). For example, histone and ribosomal proteins that are not assembled are rapidly degraded (Abovich et al., 1985; Agrawal and Bowman, 1987; elBaradi et al., 1986; Gunjan and Verreault, 2003; Maicas et al., 1988; Warner et al., 1985). Related to this, aneuploid cells are more sensitive to drugs inhibiting protein synthesis, folding, or proteasome function (Tang et al., 2011; Torres et al., 2007; Whitesell and Lindquist, 2005).

Previous studies investigating the causes of the reduction in proliferation rate in aneuploid cells were mainly conducted in disomic (*n*+1) or trisomic (2*n*+1) aneuploid cells (Dephoure et al., 2014; Oromendia and Amon, 2014; Oromendia et al., 2012; Stingele et al., 2012; Thorburn et al., 2013; Torres et al., 2010a; Torres et al., 2007; Williams et al., 2008). In *n*+1 or 2*n*+1 cells, both the balance hypothesis and the burden hypothesis predict that proliferation rate will decrease as the number of additional genes increases because more genes results in more imbalanced protein complexes. Indeed, proliferation rate is negatively correlated with the size of the additional chromosome in *n*+1 or 2*n*+1 cells (Sheltzer and Amon, 2011; Torres et al., 2007). However, tumor cells usually exhibit karyotypical abnormalities involving many chromosomes (e.g., 2*n+x* where x>1) (Holland and Cleveland, 2009; Weaver and Cleveland, 2006). 2*n+x* cells will contain a mixture of protein complexes balanced at the 2-copy level, complexes that are imbalanced, and complexes balanced at the 3-copy level. Thus, the number of imbalanced protein complexes does not necessarily monotonically increase as the number of additional genes increases in an aneuploid tumor cell. Given this emergent property, new mechanisms that underlie variation in proliferation rate in more complex aneuploid cells may exist. In this study, we used *S. cerevisiae* to generate a variety of 2*n+x* cells and determined how gene copy number affects proliferation.

## RESULTS

### Proliferation rate of a 2*n+x* yeast strain is negatively correlated with the total number of genes on these *x* chromosomes

We obtained a variety of 2*n+x* (*n*=16) budding yeast strains by collecting the meiotic products of a pentaploid strain (5*n*, **Fig. 1A** and **Fig. S1A**). We focused on 10 tetrads in which all 4 meiotic products were viable (**Fig. 1B**). Colony size varied among these strains (**Fig. 1B**). We quantified the area of each colony after a 4-day growth on the tetrad dissection plates and used this to infer the proliferation rate of these 2*n+x* strains.

**Figure 1.**
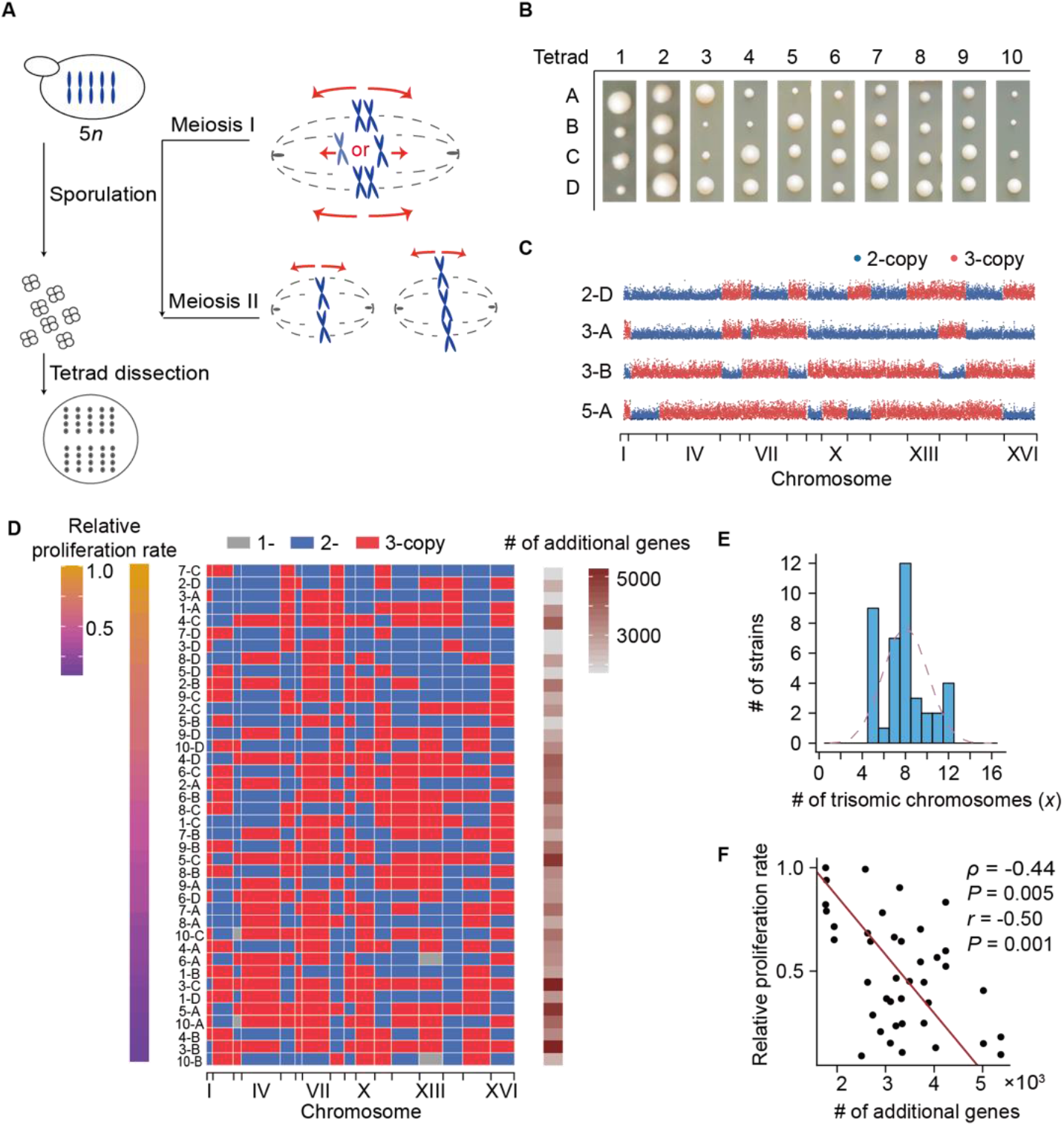
Proliferation rate is negatively correlated with the total number of additional genes. (A) A schematic description of the construction of 2*n+x* yeast strains. During meiosis I of the 5*n* strain, each chromosome randomly segregated into one of the two nuclei. (B) Colonies of 40 aneuploid strains that were derived from 10 tetrads. Four spores generated in one meiotic event are aligned in one column. (C) Scatter plots illustrating the karyotypes of four 2*n+x* strains. Dots show the average sequencing depths of non-overlapping 1000 bp windows from DNA-seq. 2-copy and 3-copy chromosomes are shown in blue and red, respectively. The 16 chromosomes are concatenated in order along the x axis. (D) Karyotypes of forty 2*n+x* strains. Strains are ordered by proliferation rate with the number of additional genes in each strain shown in the rightmost column. (E) The distribution of the number of additional chromosomes (x) among the 40 aneuploid strains. The distribution is indistinguishable from a normal distribution centered at 8 (purple dashed line, *P* = 0.61, one-sample *t* test). (F) Proliferation rate is negatively correlated with the number of additional genes. P-values were calculated with both Spearman’s correlation and Pearson’s correlation. See also Figures S1-2 and Table S1.

We performed DNA-seq to obtain the karyotypes of these strains (**Fig. 1C** and **Fig. S2**). The number of additional chromosomes per strain, x, varied between 5 and 12 among these 2*n+x* strains, with a mean of ~8 (**Fig. 1D-E** and **Table S1**). Importantly, segregation was independent among each of the 16 chromosomes (**Fig. S1B**). The total number of additional genes varied from 1,764 to 5,376 (**Fig. 1D**) and was negatively correlated with proliferation rate (**Fig. 1F**). Although this observation appears to be consistent with both the balance and the burden hypotheses and with previous findings in *n*+1 or 2*n*+1 aneuploid cells (Niwa et al., 2006; Sheltzer and Amon, 2011; Torres et al., 2007; Torres et al., 2008; Torres et al., 2010b; Williams et al., 2008), the interpretation is different, as we will explain in the following sections.

### The number of 3-copy balanced protein complexes (N_3-copy_) is negatively correlated with proliferation rate

We examined the relationship between the status of protein complexes in a 2*n+x* strain and its proliferation rate. We used a manually curated (Pu et al., 2009) set of 408 protein complexes containing 1,617 subunits. In each strain we defined a protein complex as imbalanced if some subunits of it are encoded by 2 copies of DNA whereas others are encoded by 3 copies; by contrast, a 2-copy or 3-copy balanced complex has all of its subunits encoded by 2 or 3 copies of DNA (**Fig. 2A**). We counted the number of 2-copy balanced (N_2-copy_), imbalanced (N_imb_), and 3-copy balanced (N_3-copy_) protein complexes, respectively, in each 2*n+x* strain (**Fig. 2A**). The simulation showing N_2-copy_ monotonically decreases and N_3-copy_ monotonically increases as the number of additional chromosomes increases. In contrast, N_imb_ first increases and then decreases (**Fig. 2B**). Because the total of them (N_2-copy_+N_imb_+N_3-copy_) is equal to the total number of protein complexes (408), we can draw an equilateral triangle where each dot represents a 2*n+x* strain and the distances to three edges represent the numbers of protein complexes in the above-mentioned three categories (**Fig. S3A**), for the convenience of better illustration. As expected, as the number of additional genes increased, N_2-copy_ monotonically decreased and N_3-copy_ monotonically increased. In contrast, *N*_imb_ first increased and then decreased (**Fig. 2C** and **Fig. S3A**). Therefore, the properties that affect proliferation in complex aneuploidies may be fundamentally different from those in cells that have gained or lost only a single chromosome.

**Figure 2.**
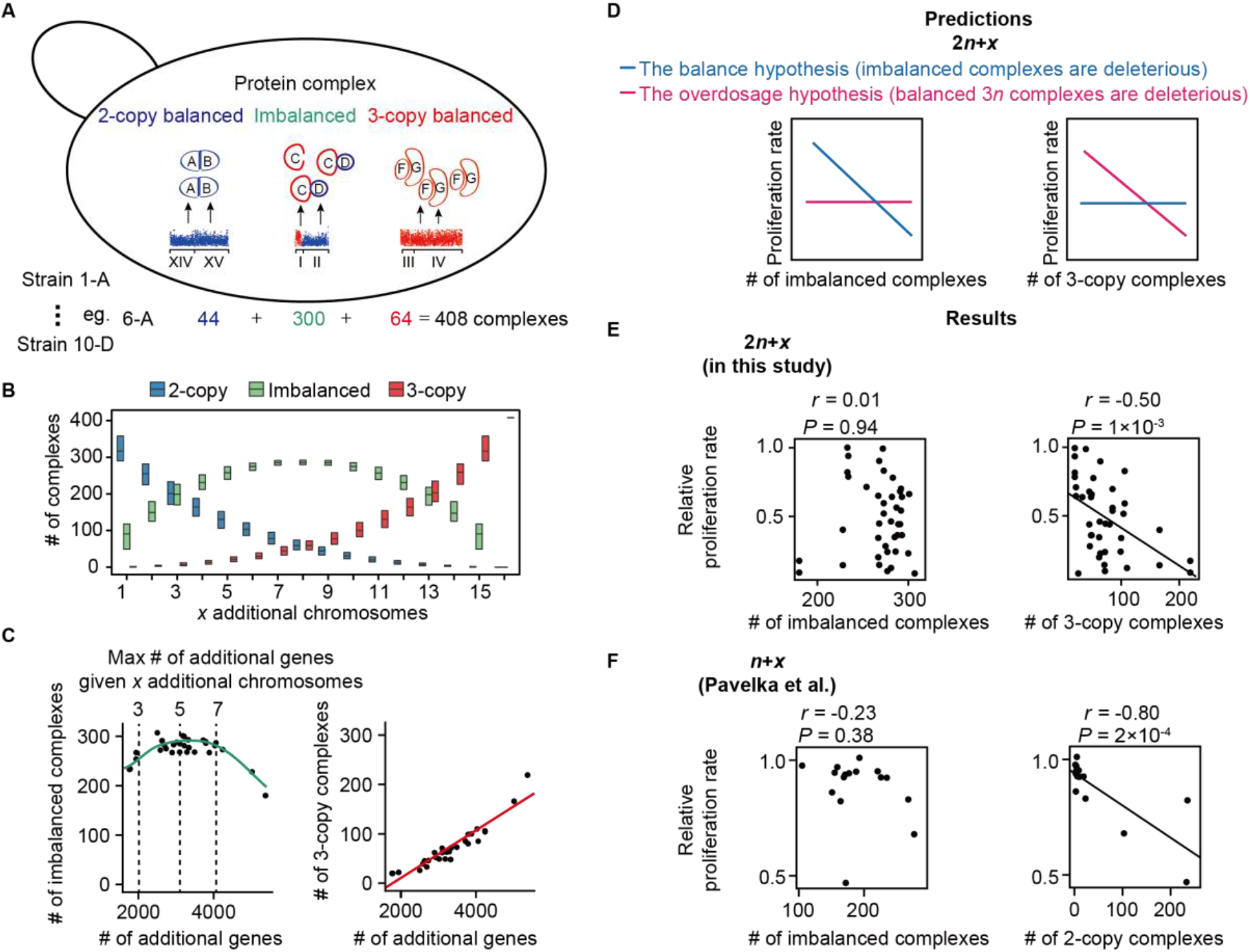
Overdosage of balanced protein complexes, but not imbalanced complexes, correlates with reduced proliferation rate in cells with complex aneuploidies. (A) In 2*n+x* strains each protein complex can be 2-copy balanced (*N*_2-copy_), imbalanced (N_imb_), or 3-copy balanced (*N*_3-copy_). The number of protein complexes in each category is shown for an example strain. (B) A simulation showing how the number of 2-copy, imbalanced, and 3-copy complexes changes with the number of additional chromosomes. Boxes show the middle 50% of 1000 simulated strains in which each strain has x random additional chromosomes. (C) The relationship between the number of additional genes in an aneuploid strain and the number of imbalanced (left) and 3-copy balanced (right) complexes. Each dot represents one 2*n+x* strain. (D) Cartoon showing the expected relation between proliferation rate and the number of additional complexes for two models for why aneuploidy causes slow proliferation. (E-F) Experimental data showing the relation between proliferation rate and the number of additional complexes in 2*n* (E) and 1n (F) strains with complex aneuploidies. See also Figures S3-4 and Table S2.

The balance hypothesis predicts a negative correlation between N_imb_ and proliferation rate (**Fig. 2D**). However, such a correlation was not observed (*r* = 0.01, *P* = 0.94, Pearson’s correlation, **Fig. 2E**). Intriguingly, the proliferation rate of 2*n+x* strains was negatively correlated with *N*_3-copy_ (*r* = −0.50, *P* = 1 × 10^−3^, **Fig. 2E**). We therefore speculated that the reduced proliferation rate in 2*n+x* strains with large numbers of 3-copy complexes was caused by an increase in dosage of entire and balanced protein complexes (the “overdosage hypothesis”) (**Fig. 2D**).

It is worth noting that the negative correlation between *N*3-copy and proliferation rate remains after controlling for confounding factors. For example, we further classified protein complexes into 20 GO categories according to their subcellular localizations and still observed a negative correlation between *N*3-copy and proliferation rate in each GO category (*P* < 0.05 in 14 categories, **Fig. S3B** and **Table S2**), suggesting that this negative relationship is independent of subcellular localizations. In addition, we separated protein complexes into two categories according to whether some subunits of the complex are encoded by genes from the whole genome duplication (WGD). The negative correlation between *N*3-copy and proliferation rate remained in both categories (**Fig. S3C**), suggesting that the overdosage of entire protein complexes reduces cell proliferation rate even among the protein complexes that are sensitive to dosage balance (Papp et al., 2003).

N_3-copy_ is highly correlated with the number of additional genes (r = 0.94, **Fig. 2C**). To determine if the correlation between *N*_3-copy_ and proliferation rate is a by-product of the correlation between the number of additional genes and proliferation rate, we further performed a permutation test to investigate the impact of *N*_3-copy_ on proliferation rate. We randomly sampled genes from the genome and assigned genes into “pseudo” protein complexes, while keeping the total number of complexes and the number of subunits in each complex unchanged. For each set of “pseudo” protein complexes we calculated the correlation between *N*_3-copy_ and proliferation rate. We performed this sampling 10,000 times and found that the experimentally observed correlation coefficient between *N*3-copy and proliferation rate was significantly stronger than the random expectation (*P* = 0.04, **Fig. S3D**).

### The number of 2-copy balanced protein complexes in *n+x* yeast strains is negatively correlated with proliferation rate

For a second test of the overdosage hypothesis, we used 16 stable *n+x* haploid yeast strains, each of which contains an extra copy of between 1 and 13 chromosomes (Pavelka et al., 2010). A 2-copy complex in an *n+x* strain (haploid background) is equivalent to a 3-copy complex in a 2*n+x* strain (diploid background) because in both cases the cell has one extra copy of complexes in addition to the 1-copy or 2-copy complement (**Fig. S4**). In agreement with the observations in 2*n+x* yeast strains, the number of two-copy balanced complexes in *n+x* strains (*N*’_2-copy_) was negatively correlated with proliferation rate (*r* = −0.80, *P* = 2×10^−4^, **Fig. 2F**), providing additional support for the overdosage hypothesis.

### Deleting one copy of a 3-copy balanced protein complex partially restored proliferation rate

To test if overdosage of a single balanced protein complex can reduce growth rate, we knocked out one copy of each gene in a 3-copy balanced protein complex and determined if proliferation rate was partially restored. To this end, we randomly chose three 2-subunit protein complexes that are balanced at the 3-copy level in a 2*n+x* aneuploid strain (see Methods for details): Cdc28p/Clb5p complex, Sgv1p/Bur2p complex, and ISW1a complex. Cdc28p/Clb5p complex regulates mitotic and meiotic cell cycle, Sgv1p/Bur2p complex is involved in the transcriptional regulation, and ISW1a complex has nucleosome-stimulated ATPase activity. For each complex, one copy of one subunit was knocked out with KanMX & GFP, and that of the other with NatMX, generating a 2-copy variant marked with GFP for the competition assay (**Fig. 3A**, left side). We co-cultured the 2-copy and 3-copy variants in rich media and used flow cytometry to measure the relative frequency of each variant over time (**Fig. 3A**).

**Figure 3.**
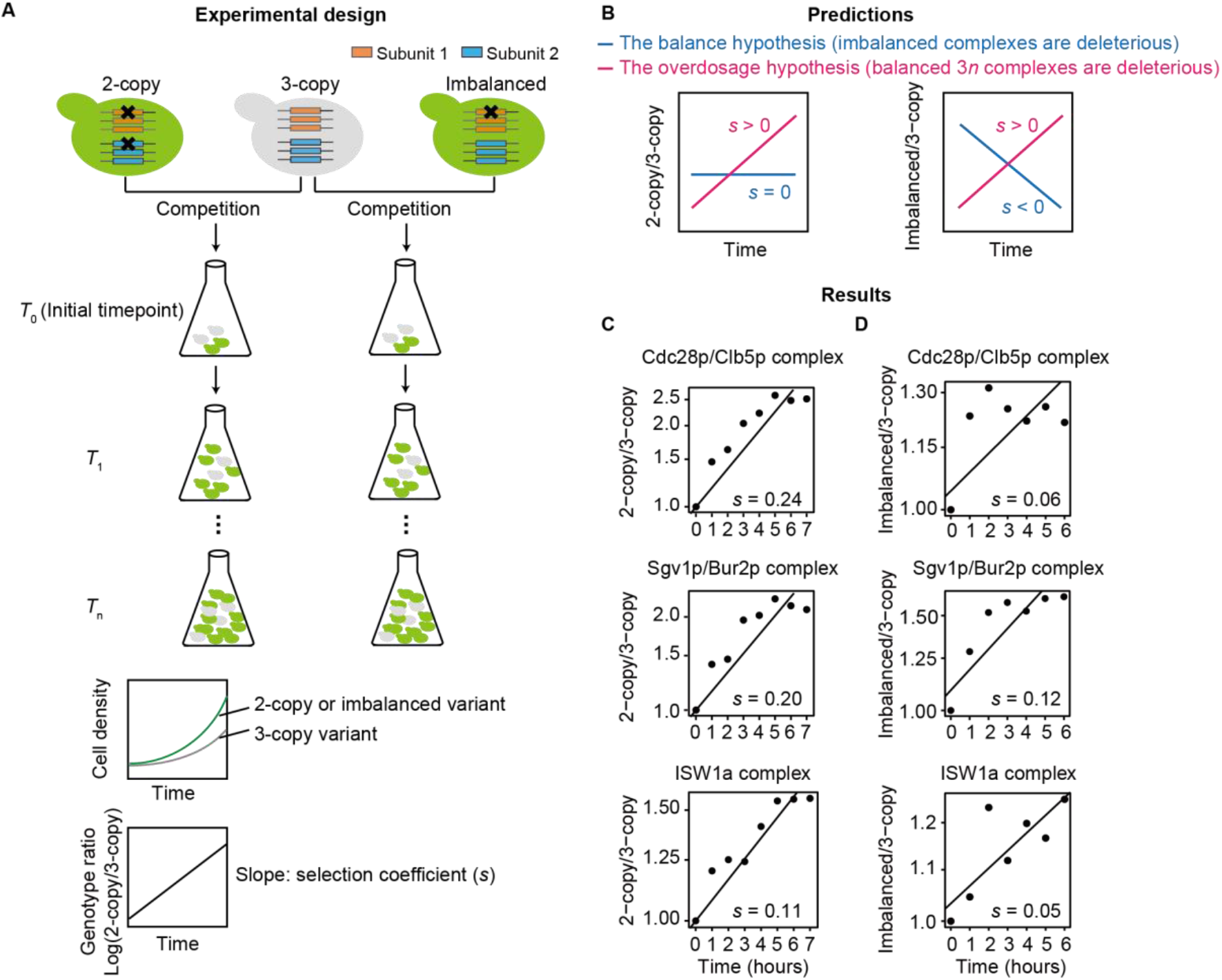
Experimental test of the balance and the overdosage hypotheses. (A) A schematic description of the construction of the 2-copy, 3-copy, and imbalanced variants, and the competition assay. The ratio of population size between 2-copy/imbalanced (GFP-positive) and 3-copy (GFP-negative) variants was determined by flow cytometry every hour. (B) Predictions of the balance and the overdosage hypotheses. (C-D) The 2-copy (C) and the imbalanced variants (D) outcompete the 3-copy variant for each of the three tested complexes.

The balance hypothesis predicts that the 2-copy and the 3-copy variants should have a similar growth rate because the copy number of each gene within the protein complex is the same (**Fig. 3B**). In contrast, the overdosage hypothesis predicts that the 2-copy variant should have a higher growth rate because it confers the wild-type concentration of the protein complexes (**Fig. 3B**). We found that the 2-copy variant outcompeted the 3-copy variant for each of the three tested complexes (**Fig. 3C**), supporting the overdosage hypothesis.

A more rigorous test to distinguish between the balance and the overdosage hypotheses is to convert a balanced 3-copy complex to an imbalanced complex by deleting a copy of only one subunit (**Fig. 3A**, right side). The balance hypothesis predicts that the imbalanced strain should have a worse proliferation rate than the 3-copy variant (**Fig. 3B**). By contrast, the overdosage hypothesis predicts that this imbalanced strain should have a higher proliferation rate than the 3-copy variant, because the concentration of the protein complex is reduced to the wild-type level (**Fig. 3B**). The imbalanced variant outcompeted the 3-copy variant for each of the three tested complexes (**Fig. 3D**), in support of the overdosage hypothesis.

### The subunits of imbalanced, but not 3-copy balanced protein complexes are degraded in aneuploid strains

Why are 3-copy balanced protein complexes deleterious? The unassembled subunits of imbalanced protein complexes are degraded (Dephoure et al., 2014; Goncalves et al., 2017; Ishikawa et al., 2017; McShane et al., 2016; Ryan et al., 2017; Stingele et al., 2012) (**Fig. 4A**). We speculated that when all subunits of a protein complex are balanced at the 3-copy level, the extra proteins are assembled and less likely to be degraded, leading to an increased protein complex dosage (**Fig. 4A**). This increase may create a stoichiometric imbalance at a higher level, e.g., within a biochemical pathway or signaling pathway, between transcription factors and their DNA targets, etc., and causes the reduction in proliferation rate.

**Figure 4.**
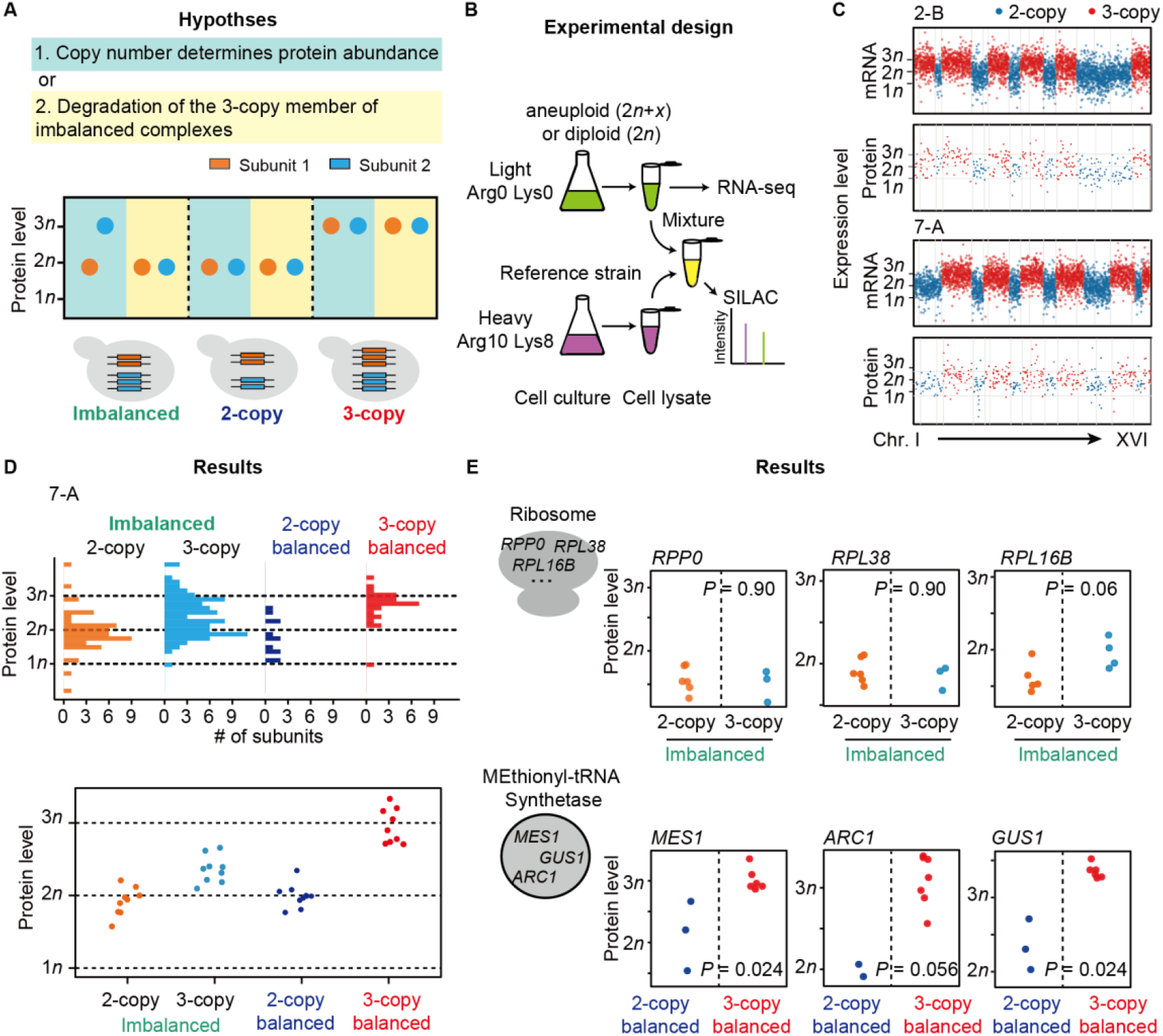
The subunits of 3-copy balanced protein complexes are not specifically degraded in *2n+x* strains. (A) The predictions of protein levels of 3-copy subunits in imbalanced and balanced protein complexes. (B) A schematic illustration of SILAC analysis on 2*n+x* cells. (C) mRNA and protein abundances largely scale with gene copy numbers. Each dot represents one gene. Blue and red dots represent 2-copy and 3-copy genes, respectively. 16 chromosomes are concatenated in order on the x axis. (D) The measured relative abundance of each protein from each strain, split into imbalanced and balanced classes. (E) Relative protein abundance of six genes from all nine strains. See also Figure S5.

To test this hypothesis we quantified the transcriptomes and proteomes in nine 2*n+x* strains and in a diploid 2*n* strain (**Fig. 4B-C**). For each 2*n+x* strain we divided genes into three classes: in an imbalanced protein complex, in a 2-copy balanced protein complex, and in a 3-copy balanced protein complex (**Fig. 4D**). In balanced protein complexes, the majority of 3-copy genes exhibited on average a 3n protein level (**Fig. 4D**), consistent with their increase in both DNA and mRNA (**Fig. S5A**). In contrast, in imbalanced protein complexes, the protein concentrations of ~50% of the 3-copy genes were more similar to those of 2-copy genes (**Fig. 4D**), although their mRNA concentrations still exhibited a 0.5-fold increase (**Fig. S5A**).

Examination of two protein complexes showed that when all subunits of a protein complex are balanced at the 3-copy level, the protein levels of these subunits increased (**Fig. 4E** and **Fig. S5B-D**). In contrast, the additional subunits in imbalanced protein complexes were specifically degraded (**Fig. 4E** and **Fig. S5B-D**). These observations suggest that the DNA copy number increase of all subunits in a protein complex facilitates the escape from surveillance mechanisms that specifically degrade individually overexpressed subunits. This escape leads to the overdosage of the protein complex.

### Dosage-sensitive protein complexes are enriched in the cell cycle pathway

To understand why the overdosage of balanced protein complexes is deleterious, we identified individual dosage-sensitive protein complexes that lead to a reduction in proliferation when present in 3 copies. For each complex, each 2*n+x* strain can be classified into one of the following three groups (**Fig. 5A**). If all subunits of the complex are at the 2-copy (or 3-copy) level, we classified this strain into group 2 (or group 3). By contrast, if some subunits are at the 2-copy level whereas others are at the 3-copy level, we classified this strain into group *i* (imbalance). The overdosage hypothesis predicts that the strains in group 3, where the dosage of balanced protein complexes is higher, will exhibit significantly lower proliferation rates than those in group 2 (**Fig. 5B** and **Fig. S4**). We compared the proliferation rates between these two groups of strains for each protein complex and in total identified 123 dosage-sensitive protein complexes (*P* < 0.05, *t* test, **Table S3** and **Fig. S6**). Tubulins as an example were shown in **Fig. 5B**.

**Figure 5.**
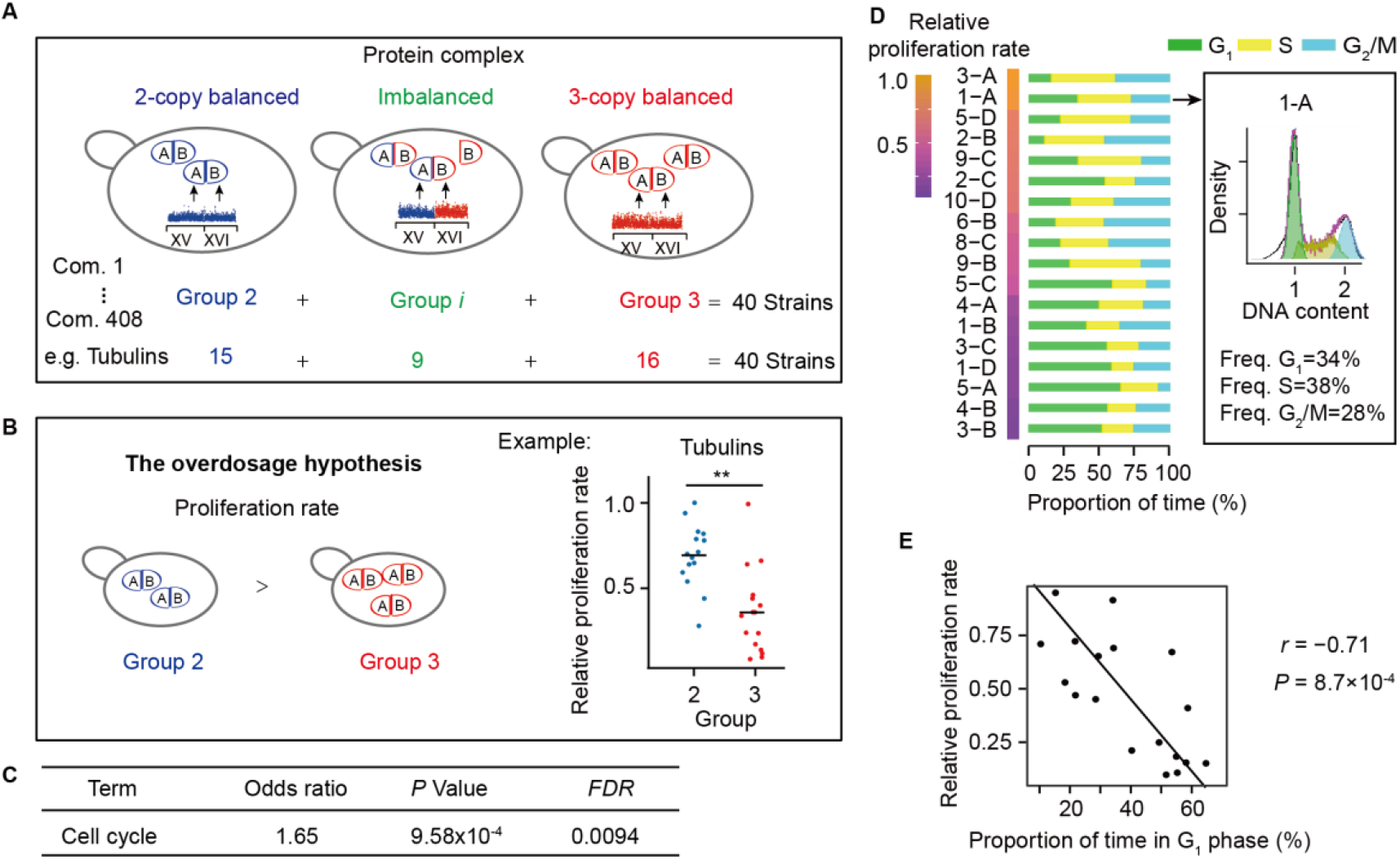
Protein complexes with support for the overdosage hypothesis are enriched in the cell-cycle pathway. (A) For each protein complex, forty 2*n+x* strains were classified into three groups (group 2, group *i*, and group 3). (B) 123 protein complexes whose dosage is negatively associated with proliferation rate are identified. Tubulins is shown as an example. (C) The results of KEGG enrichment analysis. (D) 2n+x cells exhibit delays at various stages of a cell cycle. The box shows an example how proportion of time in each cell-cycle stage was quantified. Strains are ordered by proliferation rate. (E) Proliferation rate is negatively correlated with the proportion of time in G1 stage. See also Figures S4 and S6, and Tables S3-S4.

We retrieved all 387 genes encoding the subunits of these dosage-sensitive protein complexes (**Fig. 5B**). Intriguingly, the only KEGG term where these genes were enriched was “cell cycle” (False discovery rate *FDR* = 0.01, **Fig. 5C** and **Table S4**). This is in agreement with the previous findings that cell-cycle abnormality was common among aneuploid cells (Beach et al., 2017; Thorburn et al., 2013; Torres et al., 2007). To quantify the cell-cycle defects in these strains, we used Sytox Green straining and flow cytometry to analyze the cell-cycle distribution, and observed that a cell-cycle delay in G1 stage was negatively correlated with proliferation rate (r = −0.71, *P* = 8.7× 10^−4^, **Fig. 5D-E**), suggesting that the variation in proliferation rate among our 2*n+x* strains can be partly explained by the overdosage of cell cycle-related complexes that lead to abnormality in cell-cycle progression.

### Overdosage of entire protein complexes is depleted human cancer cells

To determine if what holds for yeast cells also applies to human cancer cells, we tested the overdosage hypothesis with copy number variation data from 10,995 cancer samples across 26 cancer types collected in The Cancer Genome Atlas (TCGA) (Cancer Genome Atlas Research Network et al., 2013). We retrieved the information of 1,521 human protein complexes annotated in the Human Protein Reference Database (Keshava Prasad et al., 2009). For each protein complex and in each cancer type, we counted the number of cancer samples with this complex balanced at the 3-copy level (S3). As a control, we shuffled the DNA copy numbers of genes encoding subunits of all protein complexes within each cancer sample to obtain the expected S3. We identified 85 protein complexes in which the observed *S*3 were significantly smaller than the expected *S*3 *(P* < 0.05, paired *t* test, *df* = 25, **Fig. 6A-B, Table S5**). The subunits of these 85 complexes were enriched in the pathways inhibiting the proliferation of cancer cells. In addition to the cell-cycle pathway (e.g., apoptosis and p53, **Fig. 6C**) that were also identified in yeast, overdosage of protein complexes related to immune reaction was avoided as well.

**Figure 6.**
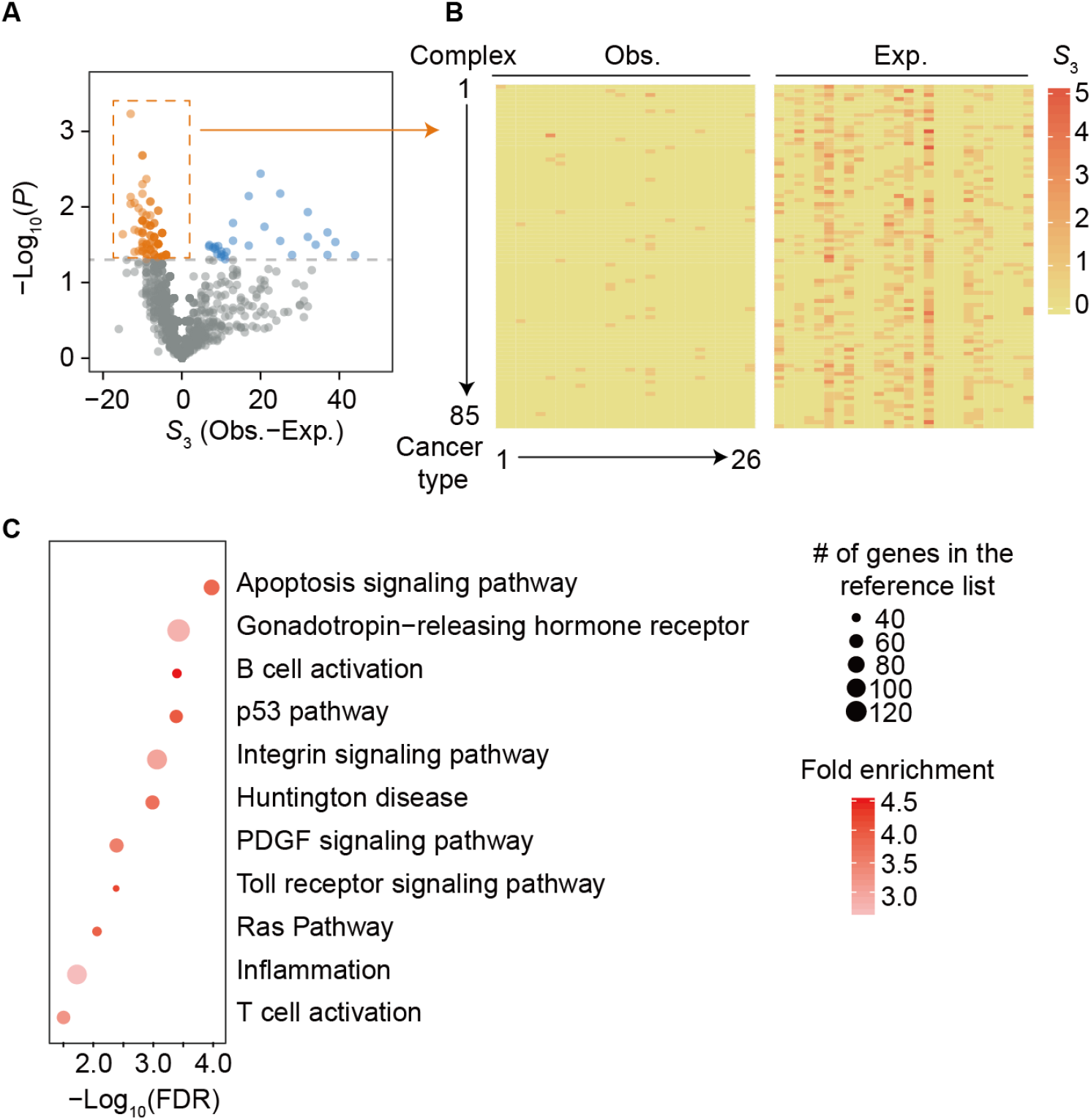
Protein complexes that are avoided to be balanced at the 3-copy level in human cancer samples. (A) Eighty-five protein complexes (dots in orange) that S_3_ are significantly smaller than the random expectation (*P* < 0.05, paired *t* test, *df* = 25) are identified. Thirty-nine protein complexes (dots in blue) that S3 are significantly greater than the random expectation are identified. Eight protein complexes exhibited a large difference (Obs.–Exp. >60) in S3 between the observation and the random expectation, because the subunits of them are encoded by genes located on the same chromosomes and are duplicated together. Therefore, the copy numbers of them tend to change coordinately. These 8 protein complexes are not shown. (B) The observed and expected S3 of the 85 protein complexes identified in (A) in each cancer type. (C) The result of PANTHER enrichment analysis on the genes encoding these 85 complexes. All genes encoding subunits of human protein complexes were used as the reference. See also Table S5.

In contrast, significantly less protein complexes exhibited greater observed *S*3 (39 vs. 85, *P* = 4.4× 10^−5^, binomial test) and these complexes were not enriched in any pathways.

## DISCUSSION

Aneuploid cells have a lower proliferation rate than euploid cells (Birchler and Veitia, 2010; Otto and Whitton, 2000), most likely due to a deviation in the balance of gene copy number (Torres et al., 2008). However, the type of genetic imbalance that causes slow growth remains elusive. Most previous studies focused on imbalance within a protein complex (Papp et al., 2003). A number of studies pointed out that aneuploidy could also lead to other types of imbalance, of the same biochemical pathway or signaling pathway, between transcription factors and their DNA targets (Birchler and Veitia, 2007, 2010, 2012; Veitia, 2004, 2005), etc. However, there was no genome-wide experimental evidence that imbalance, other than that within a protein complex, could result in a growth defect. In this study we provide the first evidence for the importance of dosage balance beyond that within a protein complex.

In this study we explicitly tested for the support of the balance hypothesis as an overall determinant of proliferation rate and for each complex independently. Whereas our data do support the balance hypothesis that copy-number imbalance of subunits within individual protein complexes results a proliferation defect (76 out of 408 protein complexes, **Table S3** and **Fig. S6**), in our data the overall effect of imbalance on proliferation rate is minimal. The balance hypothesis predicts a negative correlation between proliferation rate and N_imb_, but we did not find such correlation in either *n+x* or 2*n+x* cells.

There are at least four possible reasons why the genomic-scale imbalance (N_imb_) does not correlate with proliferation in our data. First, it is possible that severe imbalances within protein complexes were lethal, and thus, the individuals suffering from them cannot be included in this study. Second, mechanisms exist whose end result is that an increase in gene copy-number of a single subunit does not lead to an increase in protein levels (Birchler and Veitia, 2012; Dephoure et al., 2014; Ishikawa et al., 2017; McShane et al., 2016; Stingele et al., 2012). This was the dominant effect in this study; over 50% of subunits present in 3 copies in imbalanced complexes were expressed at the 2-copy level in the proteome (**Fig. 4**). A third possible reason for the lack of correlation between the number of imbalanced protein complexes (N_imb_) and proliferation rate is additional effects from other factors, namely the total number of additional genes and the number of 3-copy balanced complexes (*N*_3-copy_). Indeed, a linear model that predicts proliferation rate from all three parameters: the number of additional genes, N_imb_, and N_3-copy_, performs better (AIC = 60) than models that include the number of additional genes plus one of the other two parameters (AIC = 68.5 after removing *N*_3-copy_ and 71.7 after removing N_imb_). Therefore both the balance and the overdosage hypotheses are supported by our data. Lastly, “bridge” subunits (Bray and Lay, 1997) play a central role in the assembly of protein complexes and are likely more sensitive to stoichiometric imbalance. However, such bridge subunits have not been identified genome-widely. The effect of the imbalance of such subunits may be masked by a large number of other subunits that are not sensitive to stoichiometric imbalance.

Protein complexes that are avoided to be balanced at the 3-copy level in human cancer samples were enriched in a number of pathways (**Fig. 6**). Surprisingly, some of these signaling pathways seem to promote tumorigensis (e.g., Ras pathway). However, a careful inspection revealed that the overexpression of part of such pathways might actually suppress the proliferation of cancer cells. For example, 3 proteins (KSR1, EPHB2, and MAP2K1) in the Ras pathway form a protein complex and this complex was not balanced at the 3-copy level in any of the 22,457 cancer samples examined (*P* = 0.02, **Table S5**), likely because Kinase Suppressor of Ras (KSR1) suppresses tumorigensis through forming a protein complex with the other two proteins (Denouel-Galy et al., 1998; Yu et al., 1998). This is consistent with our hypothesis that the overdosage of an entire protein complex causes the stoichiometric imbalance at a higher level, such as in a signaling pathway. In addition, some other pathways (e.g., Huntington disease) seem to be unrelated to tumorigensis. However, patients with Huntington disease actually exhibited a lower cancer risk (Sorensen et al., 1999), probably due to the role of expanded polyglutamine in stimulating p53 pathway (Bae et al., 2005).

## AUTHOR CONTRIBUTIONS

Y. C. and W.Q. conceived the research; Y. C., S. C., K. L., X. H., T. L., and Y. W. performed the experiments, Y. C., Y. Z., and W.Q. analyzed the data; Y. C., S. W., L. B. C., and W. Q. wrote the manuscript.

## ACKNOWLEDGEMENTS

We thank Calum Maclean for providing yeast strains. We thank Xiaolei Su, Ye Tian, and Jianzhi Zhang for discussion and James A. Birchler, Xionglei He, and Jian-Rong Yang for critical comments on the manuscript. This work was supported by grants from National Natural Science Foundation of China to W. Q. (91731302). L.B.C. was supported by grants from the Ministerio de Economía y Competitividad (MINECO) (BFU2015-68351-P) and AGAUR (2014SGR0974 & 2017SGR1054) and the Unidad de Excelencia María de Maeztu, funded by the MINECO (MDM-2014-0370).

## CONFLICT OF INTEREST

The authors declare no conflict of interest.

## METHODS

### Construction of yeast strains with complex aneuploidies

To obtain 2*n+x* yeast strains, we first constructed a 5*n*–Chr III strain. Three haploid strains (Y12 background, *MAT***a** *hoΔ0*::HygMX4; M22 background, *MATα* hoΔ0::KanMX4; NCYC110 background, *MAT***a** *hoΔ0*::HygMX4) and a 2*n*–Chr III strain (UWOPS05-217.3 background, *MATa* hoΔ0::HygMX4) were used. The 2*n*–Chr III strain contains 1 copy of chromosome III and 2 copies of the other 15 chromosomes. Because the locus determining the mating type of yeast *(MAT)* is on chromosome III, this 2*n*–Chr III strain can mate with a *MATα* strain. The 5*n*–Chr III strain was constructed by two rounds of mating (**Fig. S1A**). In the first round, a 2*n* strain was generated by crossing Y12 and NCYC110 and a 3*n*–Chr III strain was generated by crossing UWPOS05-217.3 and M22. After that, *MATα* and *MAT***a** loci were knocked out in the 3*n*–Chr III and 2*n* strains, respectively. In the second round of mating, the 5*n*–Chr III strain was obtained by crossing the 2*n* and 3*n*–Chr III strains generated above. Three 5*n*–Chr III strains were independently generated. Interestingly, one of them gained a copy of Chr III and Chr VII and another lost a copy of Chr XIV. These initial strains were referred to as 5n for short hereafter and in the main text.

5*n* cells were sporulated to generate 2*n+x* yeast strains. Cells were pre-grown in liquid YPD culture (1% yeast extract, 2% peptone, and 2% dextrose) and were transferred to the pre-sporulation media (1% yeast extract, 2% peptone, and 1% potassium acetate), where cells grew for 18-24h to reach the late log phase (1 × 10^7^ to 2× 10^7^ cells/ml). Cells were harvested by centrifugation (3000g, 5 min) at room temperature and were washed twice with sterile distilled H2O. Cells were further resuspended in the sporulation media (1% potassium acetate) at a final density of 1× 10^7^ to 2× 10^7^ cells/ml and were incubated in a shaking incubator at 30°C and 200 rpm for 4 days, after which tetrad dissections were performed.

### Quantification of proliferation rate

Proliferation rates of these 2*n+x* strains were estimated by calculating the areas of colonies, after a 4-day growth on the tetrad dissection plates. The images of 40 colonies were converted to binary images with the function “im2bw” in MATLAB (level = 0.7). The area of each colony was qualified with the function “regionprops”. To control for batch effects across plates, the proliferation rate of each strain was divided by the average proliferation rate of all 2*n+x* strains on the corresponding plate. The proliferation rate of a 2*n+x* strain was further normalized by the maximal proliferation rate among all 40 aneuploid strains. Cells that separated in meiosis II usually had similar but not identical karyotypes and proliferation rates (**Fig. 1B, D-E**), probably due to sister chromatid mis-segregation or chromosome translocations during meiosis (Loidl, 1995). Therefore, they were treated as independent samples.

### Karyotyping

Genomic DNA of 40 aneuploid strains was sequenced with the Illumina Hiseq 2000 sequencing system. Paired-end reads (150bp×2) were aligned to the SGD R64-2-1 S288C reference genome (Cherry et al., 2012) with the Burrows-Wheeler Aligner (BWA v0.7.12). The single-nucleotide polymorphism (SNP) information was used to determine the copy number of each chromosome. To this end, the genomes of four parental strains (Y12, M22, NCYC110 and UWOPS05-217.3) were also sequenced. After sequencing reads were aligned to the S288C reference genome, the SNP information of these strains was obtained by SNP calling with GATK and samtools (Li, 2011; McKenna et al., 2010). Only concordant SNPs that were called by both tools were used for further analysis. In total, we identified 24751, 38179, 29333, and 33497 strain-specific SNPs that were unique in Y12, M22, NCYC110, and UWOPS05-217.3, respectively. We then divided each chromosome in a 2**n+x** strain into non-overlapping 2000 bp windows, and counted the number of strains that contributed to the DNA in each window based on their unique SNPs. If a chromosome contains more than 20 windows where SNPs are contributed by 3 strains, the chromosome was classified as a 3-copy chromosome. The classification was further manually curated. 3-copy chromosomes exhibited on average a ~1.5 fold sequencing read density of that of 2-copy chromosomes (**Fig. S2**), suggesting that the genome remained stable at least in the first few rounds of cell division after sporulation. Some chromosomes are 1-copy and some are chimeric, with part of them 2-copy and the rest 3-copy, probably due to the mis-segregation or chromosome translocation during meiosis. For such chromosomes, the copy numbers of the regions with a larger size are shown in **Fig. 1D**. It is worth noting that SNPs may potentially affect the proliferation rates of these 2*n+x* strains. Nevertheless, partial correlations between *T* and proliferation rate remained significant after controlling for the proportion of SNPs (Y22; M22; NCYC110; UWOPS05-217.3) from each parental strain in a 2*n+x* strain (r = −0.45, *P* = 0.005; *r* = −0.51 *P* = 0.001; *r* = −0.51 *P* = 0.001; *r* =-0.50 *P* = 0.001, respectively). Here, the SNP proportion of a parental strain in a 2*n+x* strain was calculated as the total number of reads matching the unique SNPs of the parental strain divided by the total number of reads covering the unique SNPs of this parental strain.

### Construction of 2-copy and 3-copy variants

We performed a manipulative experiment that genetically changed a 3-copy protein complex to 2-copy in a 2*n+x* aneuploid strain. Strain 8-C was chosen because it is the only 2**n+x** aneuploidy among 40 strains in this study that contains only one dominant selective marker (HygMX), and therefore, two additional genes can be deleted using KanMX and NatMX. Three 2-subunit protein complexes that are balanced at the 3-copy level were randomly chosen. For each complex, one copy of each subunit was knocked out by KanMX and NatMX, respectively. *GFP* was knocked in at the same time. A 3- copy strain was generated, in which two copies of *HO* were knocked out with these two markers, respectively, in order to control for the effect of selective markers on proliferation rate. *HO* is a gene required for homothallic switching and is believed to have no effect on vegetative growth (Qian et al., 2012) with the same selective markers, generating the 3-copy variant. Primers used in PCR-based gene replacement were listed in **Table S6**.

### Quantification of mRNA abundance

mRNA-seq was performed to quantify the mRNA abundance of 9 aneuploid strains. Paired-end reads (150bp×2) were aligned to the SGD R64-2-1 S288C reference genome (www.yeastgenome.org) using Tophat (v.2.1.0), allowing up to 5 mismatches. Transcript abundance, in the unit of Reads Per Kilobase per Million mapped reads (RPKM), was calculated with cufflinks (v2.2.1).

### Stable isotope labeling with amino acids in cell culture (SILAC)

A strain for heavy isotope labeling (Lys8 and Arg10) was constructed on the background of BY4742 *(MATα his3Δ1 leu2Δ0 lys2Δ0 ura3Δ0)*, for SILAC analysis. BY4742 cannot synthesize lysine because *LYS2* was knocked out. We further knocked out *ARG4* and *CAR1 (arg4Δ0::kanMX4 car1Δ0::LEU2)* so that this strain can neither synthesize arginine nor convert arginine to proline. Primers used in PCR-based gene replacement experiments were listed in **Table S6**.

Aneuploid cells were cultured overnight at 30°C in a synthetic complete medium in the presence of light amino acid isotopes (Lys0 and Arg0, 100 mg/ml each). The reference strain (BY4742 *MATα his3Δ1 leu2Δ0 lys2Δ0 ura3Δ0 arg4Δ0::kanMX4 car1Δ0::LEU2)* was cultured similarly but in the presence of heavy amino acid isotopes (^15^N2^13^C6-lysine and ^15^N4^13^C6-arginine, Lys8 and Arg10 respectively for short). Cells were harvested at the mid-log phase (OD660 = 0.6-0.8) and were resuspended in 150 μl lysis buffer (8M urea, 1 mM sodium orthovanadate, 1 mM sodium fluoride, 2.5 mM sodium pyrophosphate, 1 mM B-glycerophosphate, 0.2% tablet of protease inhibitor, and 1 mM PMSF). Subsequently, we mixed the cell lysate of each light amino acid-labeled 2*n+x* or 2*n* strain with that of the heavy amino acid-labeled reference strain under the 1:1 ratio. The mixed lysate was then sonicated for 20 min (with a basic cycle of 10 seconds on and 10 seconds off, 30% power) with an ultrasonic homogenizer (Scientz Biotechnology). After a centrifugation at 12,000 rpm for 20 min at 16°C, the supernatant was subject to the tandem mass spectrometry (MS/MS) analysis at the Proteome Core Facility at IGDB. MS/MS spectra were matched to the database containing the translated sequences of all predicted open reading frames in the *S. cerevisiae* genome (Cherry et al., 2012) and protein abundances were estimated with MaxQuant (v1.4.1.2) (Cox et al., 2009; de Godoy et al., 2008). Search parameters allowed less than 2 missed cleavages and up to 3 labeled amino acids to be detected in a peptide sequence.

### DNA content analysis with fluorescent activated cell scanning

Aneuploid cells were cultured in YPD liquid media and 5× 10^6^ cells were harvested at the log phase. A 2*n* strain generated by crossing Y12 and NCYC110 and a 3*n* strain generated by crossing the 2*n* strain and M22 were harvested in parallel as the diploid and triploid reference genome. Cells were fixed with 200 μl 70% ethanol for 1 h at room temperature, and were subsequently incubated in 200 μl 50 μg/ml RNase A solution for 3 h at 37°C and 200 μl 50 μg/ml proteinase K solution for 3 hours at 55°C. Both reagents were dissolved in 0.05 M sodium citrate. Samples were sonicated using an ultrasonic homogenizer (60% power) for 2 cycles (10 seconds on and 10 seconds off) to disperse cells evenly. Cells were stained with Sytox Green (Life Technologies) for 20 minutes following the manufacturer’s instructions. Finally, DNA content was analyzed with flow cytometry (BD FACSAria II, excitation at 488 nm). The proportion of time in each cell-cycle stage was qualified by fitting the cell-cycle model with FlowJo.

### Data retrieval

An up-to-date set of protein complexes in yeast (CYC2008 complexes set) was downloaded from http://wodaklab.org/cyc2008/downloads (Pu et al., 2009). Subunits encoded by genes on the mitochondrial genome were not included in this study. Genes from the WGD were retrieved from a previous study (Kellis et al., 2004). The karyotypes and proliferation rates of *n+x* aneuploid strains (in rich media at 23°C) were retrieved from the study by Pavelka et al and were downloaded from the Open Data Repository (ODR) of the Stowers Institute for Medical Research (http://odr.stowers.org/websimr/). The data of copy number variation in cancer cells were downloaded from The Cancer Genome Atlas (TCGA) database (Cancer Genome Atlas Research Network et al., 2013), which includes the information of 22,457 human cancer samples across 26 cancer types. The information of 1,521 protein complexes in humans was downloaded from HPRD release 9 (www.hprd.org). 1,317 of them were annotated completely and were used in this study. Chromosomal locations of these genes were retrieved from Ensembl release 87 (www.ensembl.org).

### Data availability

The raw high-throughput sequencing data reported in this study have been deposited to the Genome Sequence Archive (Wang et al., 2017) in BIG Data Center (http://bigd.big.ac.cn/gsa), Beijing Institute of Genomics, Chinese Academy of Sciences, under accession number CRA000490. The mass spectrometry proteomics data have been deposited to the ProteomeXchange Consortium (http://proteomecentral.proteomexchange.org) via the PRIDE partner repository (Vizcaino et al., 2013) with the dataset identifier PXD007157.

